# Humoral and T-cell-mediated responses to a pre-clinical Zika vaccine candidate that utilizes a unique insect-specific flavivirus platform

**DOI:** 10.1101/2023.03.01.530296

**Authors:** Danielle L. Porier, Awadalkareem Adam, Lin Kang, Pawel Michalak, Juselyn Tupik, Matthew A. Santos, Christy Lee, Irving C. Allen, Tian Wang, Albert J. Auguste

## Abstract

Vaccination is critical for the control and prevention of viral outbreaks, yet conventional vaccine platforms may involve trade-offs between immunogenicity and safety. Insect-specific viruses have emerged as a novel vaccine platform to overcome this challenge. Detailed studies of humoral and T-cell responses induced by new insect-specific flavivirus (ISFV)-based vaccine platforms are needed to better understand correlates of protection and improve vaccine efficacy. Previously, we used a novel ISFV called Aripo virus (ARPV) to create a Zika virus (ZIKV) vaccine candidate (designated ARPV/ZIKV). ARPV/ZIKV demonstrated exceptional safety and single-dose efficacy, completely protecting mice from a lethal ZIKV challenge. Here, we explore the development of immune responses induced by ARPV/ZIKV immunization and evaluate its correlates of protection. Passive transfer of ARPV/ZIKV-induced immune sera to naïve mice prior to challenge emphasized the importance of neutralizing antibodies as a correlate of protection. Depletion of T-cells in vaccinated mice and adoptive transfer of ARPV/ZIKV-primed T-cells to naïve mice prior to challenge indicated that ARPV/ZIKV-induced CD4^+^ and CD8^+^ T-cell responses contribute to the observed protection but may not be essential for protection during ZIKV challenge. However, vaccination of Rag1 KO, Tcra KO, and muMt^−^ mice demonstrated the critical role for ARPV/ZIKV-induced T-cells in developing protective immune responses following vaccination. Overall, both humoral and T-cell-mediated responses induced by ISFV-based vaccines are important for comprehensive immunity, and ISFV platforms continue to be a promising method for future vaccine development.

## 1. Introduction

The *Flavivirus* genus has a near-global distribution and remains a significant threat to human and animal health. Flaviviruses continue to emerge worldwide, causing significant morbidity and mortality [1-4]. In particular, Zika virus (ZIKV) caused smaller outbreaks throughout the South Pacific until 1997 [5,6], before causing an explosive epidemic in Central and South America that resulted in over half a million suspected cases in South America between 2015 and 2017 [7]. The virus has caused an average loss of over 44,000 disability-adjusted life years (DALY) per year between 2010 and 2019 [8].

Despite the global health burden of ZIKV, licensed vaccines for humans remain elusive. Currently, there are only two candidates in active Phase II clinical trials: NCT NCT04917861 developed by Moderna and NCT05469802 developed by Takeda, but none have yet moved to Phase III clinical trials or been approved for use in humans. Previously, we created a recombinant Zika vaccine candidate (called ARPV/ZIKV) comprised of the genes for key antigenic Zika proteins (precursor membrane (PrM) and envelope (E)) on an Aripo virus (ARPV) backbone [9]. ARPV is a recently discovered insect-specific flavivirus (ISFV) that was isolated from *Psorophora albipes* mosquitoes in Trinidad [10]. Insect-specific viruses are incapable of replication within vertebrate hosts, and within the past decade they have emerged as a promising tool for controlling vertebrate-pathogenic viruses. They have been used to create several new vaccine candidates for both alphaviruses and flaviviruses [9,11-16]. These “pseudo-inactivated” recombinant vaccines are extremely safe due to the natural vertebrate host-restriction conferred to them by the insect-specific virus backbone, yet still retain significant immunogenicity. ARPV/ZIKV is non-pathogenic in vertebrates and cannot revert to a pathogenic form due to the vaccine’s replication deficiency in vertebrate cells [9]. Yet, a single dose of unadjuvanted ARPV/ZIKV completely protected mice from weight loss, death, viremia, and *in utero* transmission after ZIKV challenge [9]. ARPV/ZIKV’s high degree of efficacy is, in part, attributable to its rapid and robust induction of high neutralizing antibody (nAb) titers [9].

During ZIKV infection, humoral nAb responses are critical for controlling viremia and mediating protection [17,18]. CD4^+^ T-cells are also important for host protection against ZIKV-induced disease, especially for their role in generating ZIKV-specific humoral responses [19], but may also play a limited role in ZIKV clearance [19,20]. Murine and non-human primates studies have also demonstrated a role for CD8^+^ T-cell responses in protecting against viral dissemination and persistence in tissues, even though they may not be critical for protection [18,20-22]. Cross-reactivity among flavivirus antibodies at sub-neutralizing levels also raises concerns for antibody-dependent enhancement of disease (ADE) [17]. However, robust T-cell responses may help alleviate concerns about inducing DENV ADE after ZIKV vaccination [17,18].

Insect specific virus-based vaccines, especially ISFV-based vaccines, are a relatively new field of study, and a better understanding of their correlates of protection is required. Optimizing vaccine-induced T-cell responses (in addition to nAb titers) is likely critical for effective ISFV-based vaccines, such as ARPV/ZIKV. However, the role of T-cell and nAb responses in ARPV/ZIKV-induced protection against Zika disease is unknown. Here, we use ARPV/ZIKV as a model to examine the roles of humoral and cell-mediated adaptive immunity induced by ISFV-based vaccines in murine models. Through the use of passive transfer, adoptive transfer, and T-cell-depletion studies in mice, we found that ARPV/ZIKV-induced protection is primarily mediated by nAb during challenge. However, by vaccinating various genetic knockout (KO) mice, we showed that ARPV/ZIKV-induced T-cell responses are critical for the development of immunity post-vaccination. Overall, these data indicate that ISFV-based vaccines may benefit from optimization strategies to boost T-cell effector functions that are independent of their influence on nAb development.

## 2. Materials and Methods

### 2.1. Cell Lines, Viruses, and Quantification

VERO 76 and C6/36 cells from ATCC (Manassas, VA, USA) were maintained according to ATCC guidelines. ARPV isolation and ARPV/ZIKV development was previously described [9,10]. All viruses were maintained in C6/36 cells. Viral RNA was extracted using QIAmp Viral RNA Mini kits (QIAGEN, Venlo, Netherlands), according to QIAGEN instructions. Viral quantification was performed using reverse transcription quantitative PCR (RT-qPCR) (described [9]) or by plaque assay on VERO 76 cells. ZIKV-specific nAb titers in blood were quantified by plaque reduction neutralization test (PRNT_50_ or PRNT_80_) on VERO 76 cells as previously described [23], but incubations were maintained for 4 days before fixation.

### 2.2. RNA-Seq of Differentially Expressed Host Genes

Bone-marrow derived monocytes from C57BL/6J mice were differentiated into macrophages *in vitro* as previously described [10]. Macrophages were infected in triplicate with either ARPV/ZIKV, UV-inactivated ARPV/ZIKV, or ZIKV DakAr D 41524, or mock-infected with culture media. Six hours post-infection, RNA was extracted, RNA-seq conducted, and analysis performed as previously described [10] to compare the transcriptomes and identify up- or down-regulation of host innate immunity pathways.

### 2.3. Animal Experiments

All experimental animal procedures were approved by Virginia Tech Institutional Animal Care and Use Committee. ARPV/ZIKV and ARPV inoculums were prepared as previously described [9] and diluted with phosphate buffered saline (PBS) to achieve the desired dose. All mouse strains were purchased from Jackson Laboratory (Bar Harbor, ME, USA). IFN-αβR^-/-^ mice (strain #032045-JAX) were subsequently bred in-house. Four-week-old mice were divided randomly into groups prior to subcutaneous (s.c.) vaccination with 10^12^ genome copies (GC) of ARPV/ZIKV, immunization with 10^10^ GC ARPV (vaccine backbone control) or 10^8^ GC of mouse-attenuated ZIKV PRVABC59 (immunogenic positive control), or sham-vaccinated with PBS (naïve control). At 26 or 28 days post-vaccination (dpv), blood was collected retro-orbitally. The presence of ZIKV-specific nAb in serum prior to challenge was assessed by PRNT.

A lethal dose of low-passage ZIKV DakAr D 41524 (10^5^ plaque-forming units (pfu); 10^6^ GC) was administered s.c. at challenge. In the studies below, healthy control mice (unchallenged group) that were initially immunized with PBS were administered PBS rather than ZIKV at challenge. Sham-vaccinated control mice were initially immunized with PBS-diluent and received the full ZIKV challenge dose. Mice were monitored daily for clinical signs of disease post-challenge. Blood was collected for up to 4 days post-challenge (dpc) to quantify viremia. Mice were euthanized after losing 20% or more of their original body weight, or upon demonstrating severe clinical symptoms such as lethargy, hunched posture, paralysis, or unresponsiveness.

### 2.4. Passive Transfer

C57BL/6J mice (strain #000664) were vaccinated as described above. Blood was collected 28 dpv for evaluation of ZIKV nAb titers as described above. At 37 dpv, mice (n = 10-15) were euthanized and sera collected via cardiac puncture. Sera were pooled according to immunization group and ZIKV-specific nAb titers were assessed by PRNT for each pool. Pooled serum was transferred via intraperitoneal (i.p.) injection into 8-9-week-old mixed gender IFN-αβR^-/-^ mice (n = 7; 340 μL serum per mouse). Sham and unchallenged IFN-αβR^-/-^ mice received undiluted serum from mice that were initially immunized with PBS. IFN-αβR^-/-^ mice in the ARPV group and ZIKV group received undiluted serum from ARPV-immunized or ZIKV-immunized mice, respectively. Other IFN-αβR^-/-^ mice received ARPV/ZIKV serum in one of three different doses: (1) undiluted ARPV/ZIKV immune serum (high dose), (2) ARPV/ZIKV immune serum diluted 1:2 with PBS just prior to transfer (medium dose), or (3) ARPV/ZIKV immune serum diluted 1:5 (low dose). One day after transfer, sera were collected from IFN-αβR^-/-^ mice to quantify circulating nAb titers post-transfer. Two days post-transfer, IFN-αβR^-/-^ mice were challenged as described above, except for the unchallenged group which received PBS at challenge. Mice were monitored and viremia determined as described above.

### 2.5. T-Cell Depletion

Mixed gender IFN-αβR^-/-^ or C57BL/6J mice were vaccinated and challenged as described above. Monoclonal antibodies were purchased from Bio X Cell (Lebanon, NH, USA): anti-CD4 (clone GK1.5), anti-CD8α (clone 2.43), and IgG2b isotype control (clone LTF-2). On -3, -1, +1, +3, and +6 dpc, mice in each immunization group received either 100 μg anti-CD4^+^ antibody, 250 μg anti-CD8^+^ antibody, both, 100 μg isotype antibody, or an equivalent volume of PBS (no depletion group) by i.p. injection (n = 4-7). C57BL/6J mice were also administered anti-mouse IFNAR-1 monoclonal antibody (clone MAR1-5A3) (Leinco Technologies, Fenton, MO, USA) to transiently inhibit Type I IFN (IFN-α/β) at the time of ZIKV challenge, as previously described [9]. Mice were challenged 30 dpv (i.e., 0 dpc). A group of age-matched, mixed gender mice (n = 2 per depletion type) were euthanized immediately prior to challenge and bled to confirm the efficiency of T-cell depletion. Blood was collected in BD Microtainer™ K_2_EDTA tubes (BD Biosciences, San Jose, CA, USA).

Lymphopure density gradient medium (BioLegend, San Diego, CA, USA) was used to separate lymphocytes as previously described [24]. Cells were resuspended in 3 mL TheraPEAK™ ACK Lysing Buffer (1X) (Lonza, Basel, Switzerland). After a 2 minute incubation, 10 mL PBS was added. Cells were re-suspended in flow cytometry buffer and incubated with TruStain FeX™ PLUS anti-mouse CD16/32 (BioLegend) (0.25 μg per 10^6^ cells in 100 μL) for 5 minutes on ice. Cells were stained with either BD Pharmingen™ APC Rat Anti-Mouse CD8α antibody or BD Horizon™ BV605 Rat Anti-Mouse CD4 antibody (BD Biosciences), at a final concentration of 1:1000 or 1:200 respectively. T-cell populations were measured on a FACSAria Fusion flow cytometer. After confirming depletion of CD8^+^ or CD4^+^ T-cells via flow cytometry, mice were challenged and disease outcomes measured as described above.

### 2.6. Adoptive Transfer

C57BL/6J mice were vaccinated as described above. Spleens were harvested from mice 30 dpv and processed to collect live splenocytes in a single cell suspension. Single-cell suspensions of CD4^+^ or CD8^+^ T-cells were prepared from spleens by negative selection using magnetic beads (Miltenyi Biotec, Bergisch Gladbach, Germany). Naive 6-week-old IFN-αβR^-/-^ mice (n = 4-9) received 5 × 10^6^ cells (i.p) 24 hours prior to challenge. The mock group received T-cells isolated from naïve mice. The “no transfer” group received an equivalent volume of PBS. Mice were challenged and disease outcomes measured as described above.

### 2.7. Vaccination of Tcra KO, Rag1 KO, and muMt^−^ Mice

Rag1 KO mice (strain #002216) lack mature B-cells or T-cells, muMt^−^ mice (strain #002288) produce no mature B-cells via the *mu* gene and do not express membrane-bound IgM, and Tcra KO mice (strain #002116) have dysfunctional α/β T-cell receptors and their thymus lacks CD4^+^CD8^−^ and CD4^−^CD8^+^ T-cells. Tcra KO, Rag1 KO, and muMt^−^ mice were vaccinated (n = 6), challenged 30 dpv, monitored, and assessed for disease as described above. Anti-mouse IFNAR-1 monoclonal antibody (clone MAR1-5A3) (Leinco Technologies) was used to transiently deplete Type I IFN during challenge as previously described [9].

### 2.8. Statistical Analysis

GraphPad Prism (v9.0) was used for analyses. Data distribution and variance were evaluated for normality, and normalized by log10 transformation as needed. Log-rank (Mantel–Cox) tests were performed with Kaplan–Meier survival curves to evaluate statistical significance. PRNT and weight change data were evaluated for significance by one-way ANOVA at each day, and viremia data were evaluated by two-way ANOVA. Multiple comparisons among groups were performed ad hoc using Tukey’s test. Data in figures represent mean ± standard deviation (SD). Statistical results are noted as not significant (ns), *p* ≤ 0.033 (*), *p* ≤ 0.002 (**), *p* ≤ 0.0002 (***), or *p* ≤ 0.0001 (****).

## 3. Results

### ARPV/ZIKV infection induces gene expression of T-cell-related cytokines

Gene expression after *in vitro* infection of murine macrophages from C57BL/6J mice was analyzed using Ingenuity Pathway Analysis (IPA) as previously described [25-27]. The number of differentially expressed genes between groups are shown as volcano plots (Fig. 1a). Correlation of replicates was >0.95 in all cases, showing strong consistency. ARPV/ZIKV infection resulted in significantly increased gene expression associated with IFN-γ, NF-κB, and TLR signaling (Fig. 1b). This appeared to be primarily associated with robust TLR signaling and, intriguingly, the activation of noncanonical NF-κB signaling. Pathways associated with PRR signaling, antigen presentation, and T-cell polarization were significantly up-regulated following ARPV/ZIKV infection (Fig. 1b). Again, this appeared to be predominately driven by TLR7 signaling and increased dendritic cell function. Unexpectedly, ARPV/ZIKV infection resulted in a significant decrease in signaling pathways associated with the type-I interferon response (Fig. 1b). Further studies are needed to confirm this at a protein expression level.

**Figure 1:**
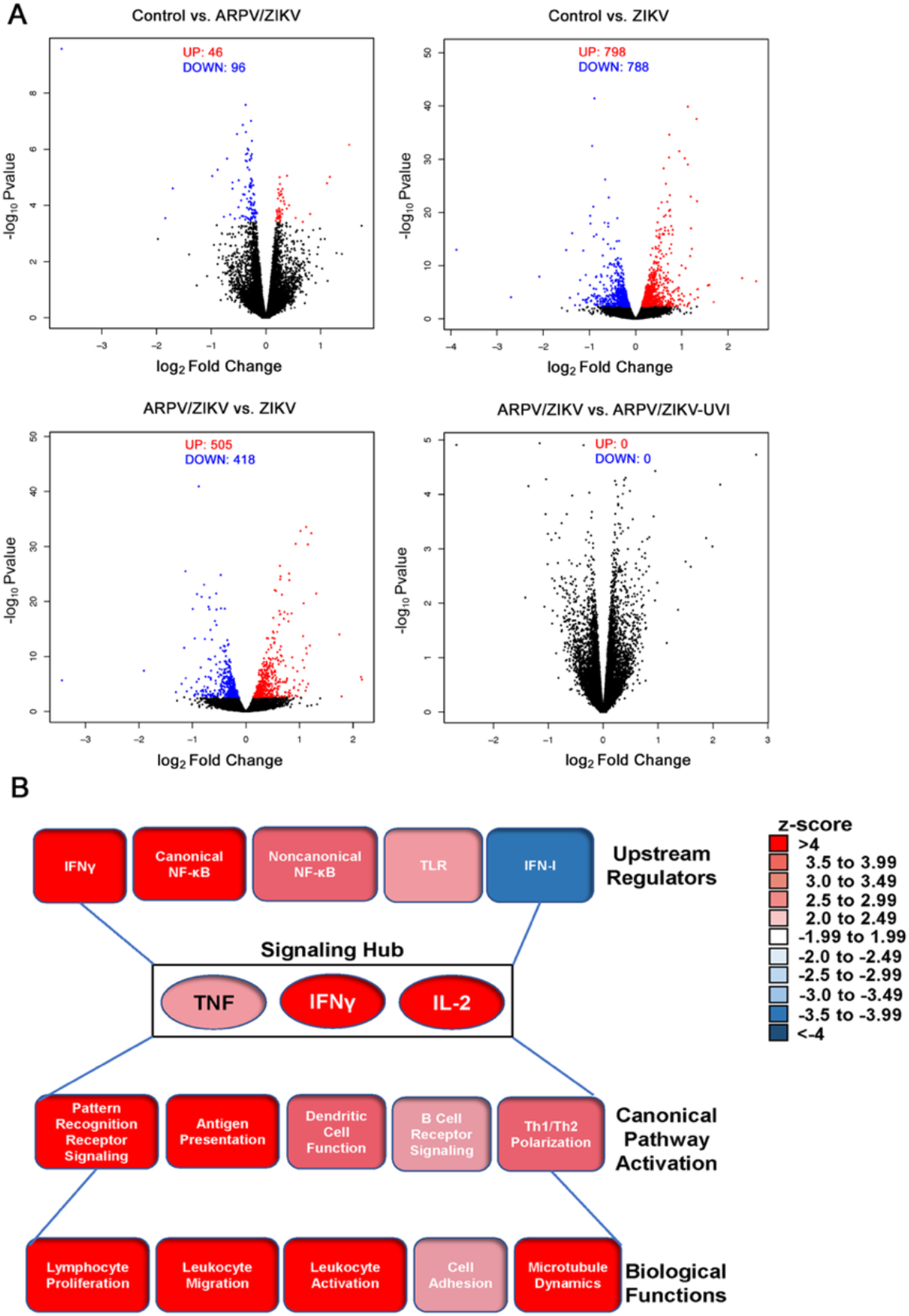
Signaling pathways associated with TNF, IFNγ, and IL-2 are significantly up-regulated by ARPV/ZIKV vaccination. Macrophages derived from naïve C57BL/6J mice were inoculated with live virus, UV-inactivated (UVI) virus, or culture media as a negative control. Host cell gene expression data was analyzed six hours post-infection, generating (a) volcano plots of differential gene expression and (b) a heatmap schematic of host immune pathways significantly up- or down-regulated after exposure to ARPV/ZIKV. The z-score range for each pathway is reflected by color intensity (up-regulation in red, down-regulation in blue). The top five pathways identified were grouped and displayed to show upstream regulators, activated canonical pathways, and predicted biological functions.

### Passive transfer of ARPV/ZIKV-induced neutralizing antibodies were unable to provide complete protection from a lethal ZIKV challenge

To determine if ARPV/ZIKV-induced nAbs responses were alone sufficient to protect mice from ZIKV, immune sera was transferred to naïve mice pre-challenge. One day pre-challenge, naïve IFN-αβR^-/-^ mice that received a high, medium, or low dose of ARPV/ZIKV immune serum had circulating ZIKV-specific nAb titers of >640 PRNT_50_ (240 PRNT_80_), 365 PRNT_50_, and 200 PRNT_50_, respectively. Mice that received undiluted ZIKV immune serum had titers of 308 PRNT_50_ (Fig. 2a). Mice that received undiluted ZIKV immune serum had the least mortality, weight loss, and viremia (Fig. 2b-d). Mice that received sera from ARPV-immunized or sham-vaccinated mice or a low dose of ARPVZIKV immune serum experienced 100% mortality by 8 dpc whereas mice that received a high dose of ARPV/ZIKV serum or ZIKV serum experienced 43% mortality by 14 dpc (Fig. 2b). Despite this similarity in overall mortality, mice that received a high dose ARPV/ZIKV immune serum demonstrated significantly more weight loss 8 dpc than mice that received ZIKV immune serum (Fig. 2c), even though ARPV/ZIKV high dose mice had higher nAb titers (Fig. 2a).

**Figure 2:**
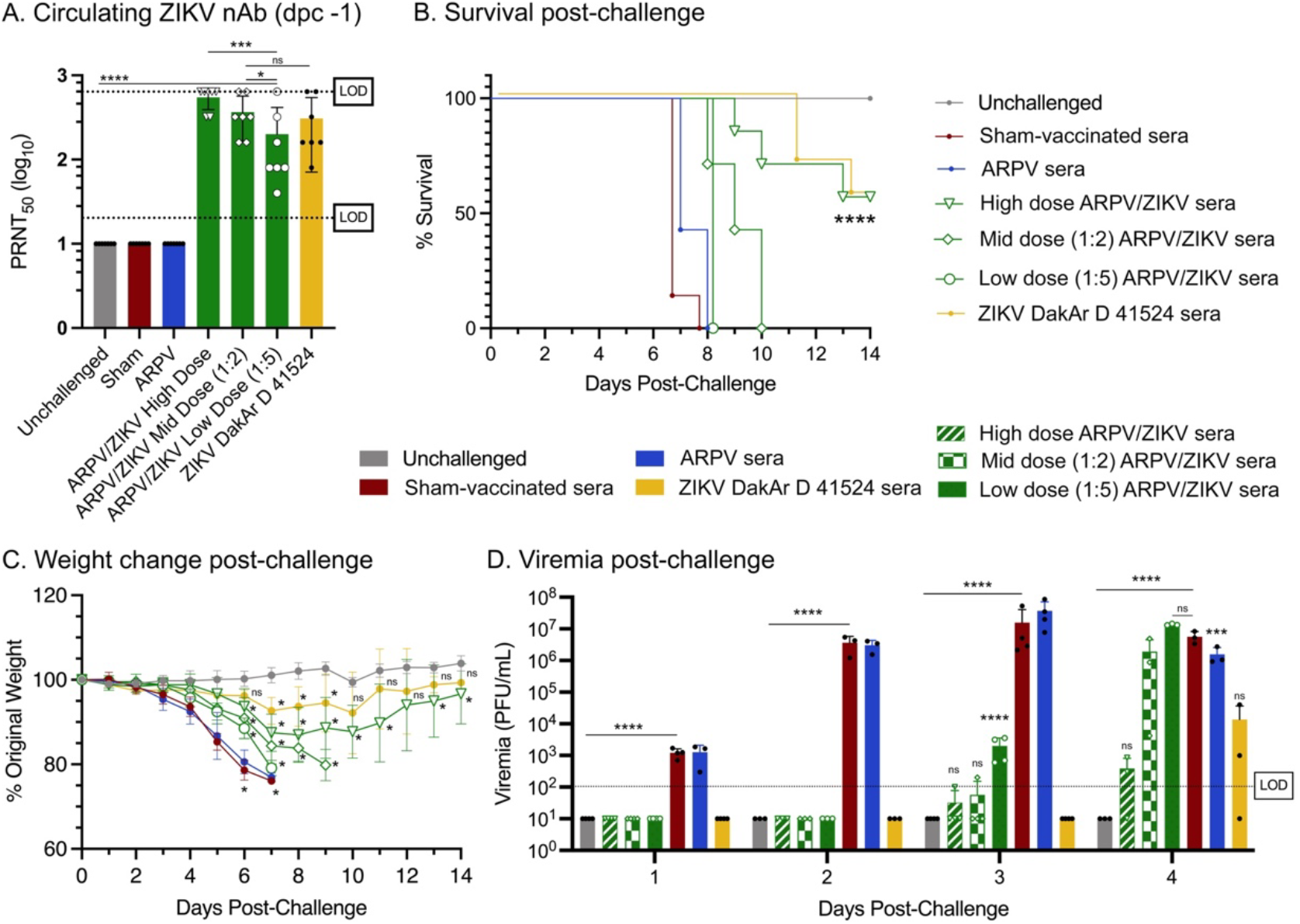
Passive transfer of ARPV/ZIKV-induced antibodies partially protects naïve mice. Four-week-old C57BL/6J mice were vaccinated as previously described. At 30 days post-vaccination, sera was pooled according to immunization group and injected into naïve 8-9 week-old IFN-αβR^-/-^ mice. (a) One day prior to challenge, mice were bled to quantify the amount of ZIKV-specific nAb circulating in the blood. Dotted lines indicate the 20-fold and 640-fold limits of detection (LOD). Mice were challenged with ZIKV, except for the unchallenged group. Mice were then monitored for (b) survival and (c) weight change post-challenge. (d) Sera was collected for 1-4 days post-challenge (dpc) to quantify viremia. LOD is 100 pfu/mL. (a, d) Columns represent mean values, and symbols represent individual data points. (c) Symbols represent mean values. Error bars indicate SD of the mean. Asterisks indicate significance compared to healthy mice (unchallenged controls), unless otherwise indicated: not significant (ns), *p* ≤ 0.033 (*), *p* ≤ 0.0002 (***), *p* ≤ 0.0001 (****)

### CD4^+^ and CD8^+^ T-cells induced by ARPV/ZIKV vaccination were not critical for protection after ZIKV challenge

IFN-αβR^-/-^ mice that were passively immunized with immune sera demonstrated significantly reduced circulating nAb titers compared to mice that were directly immunized previously [9]. To determine if the incomplete protection observed after passive transfer of ARPV/ZIKV immune sera might be due to the absence of primed T-cell responses during challenge, as opposed to reduced circulating nAb titers, vaccinated IFN-αβR^-/-^ mice (nAb titers post-vaccination shown in Fig. S1) were treated with anti-CD4^+^ and/or anti-CD8^+^ depleting antibodies immediately prior to challenge. There were no significant differences in survival, weight loss, or viremia between depletion treatments within ARPV/ZIKV-vaccinated or ZIKV-immunized mice (Fig. 3), except for the ZIKV-immunized anti-CD4^+^ and -CD8^+^ group at 6pi (Fig. 3f). Unchallenged and ARPV-immunized mice showed no significant differences in survival, weight loss, or viremia among their respective depletion groups (Fig. S2). Results depicted in Fig. 3 and Fig. S2 were also analyzed and presented according to T-cell depletion regimen rather than immunization type (Fig. S3). This depletion study was also repeated in C57BL/6J mice, although we again observed that depletion of CD4^+^ and CD8^+^ T-cells had no impact on the clinical disease experienced by ARPV/ZIKV-vaccinated mice post-challenge (Fig. 4). Confirmation of T-cell depletion in both experiments was confirmed by flow cytometry (Fig. S4).

**Figure 3:**
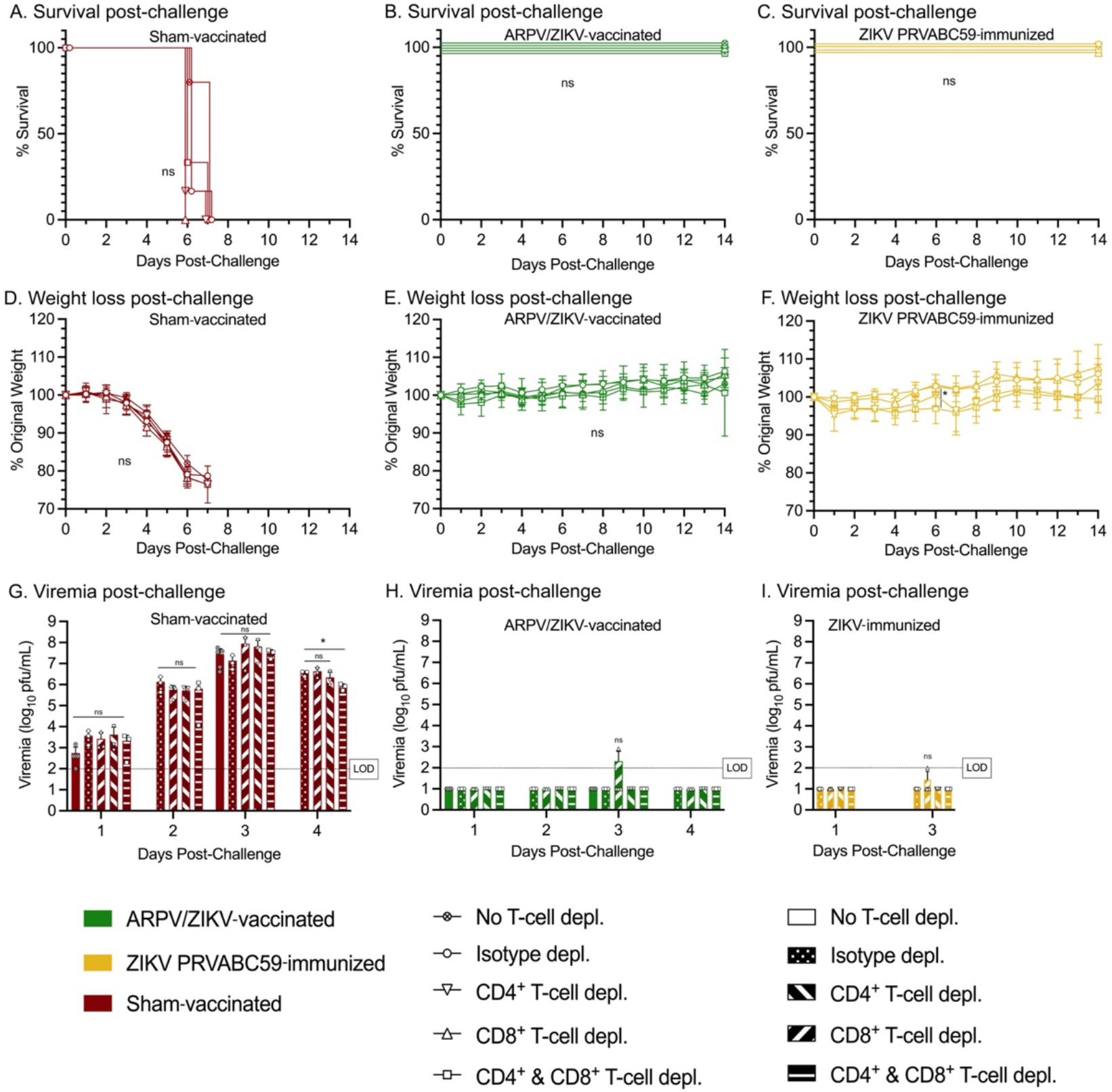
Depletion of T-cells in ARPV/ZIKV-vaccinated mice at challenge show neutralizing antibodies offer complete protection from ZIKV challenge. Four-week-old IFN-αβR^-/-^ mice were vaccinated as previously described. Beginning 27 days post-vaccination (dpv), T-cell depleting antibodies were administered according to depletion group (anti-CD4^+^, anti-CD8^+^, both, or isotype depletion). Mice were challenged with ZIKV at 30 dpv. Mice were monitored for (a-c) survival and (d-f) weight change post-challenge. (g-i) Sera was collected for 1-4 days post-challenge to quantify viremia. LOD is 100 pfu/mL. (d-f) Symbols represent mean values. (g-i) Columns represent mean values, and symbols represent individual data points. Error bars indicate SD of the mean. Asterisks indicate significance compared to isotype depleted control mice, unless otherwise indicated: not significant (ns), *p* ≤ 0.033 (*).

**Figure 4:**
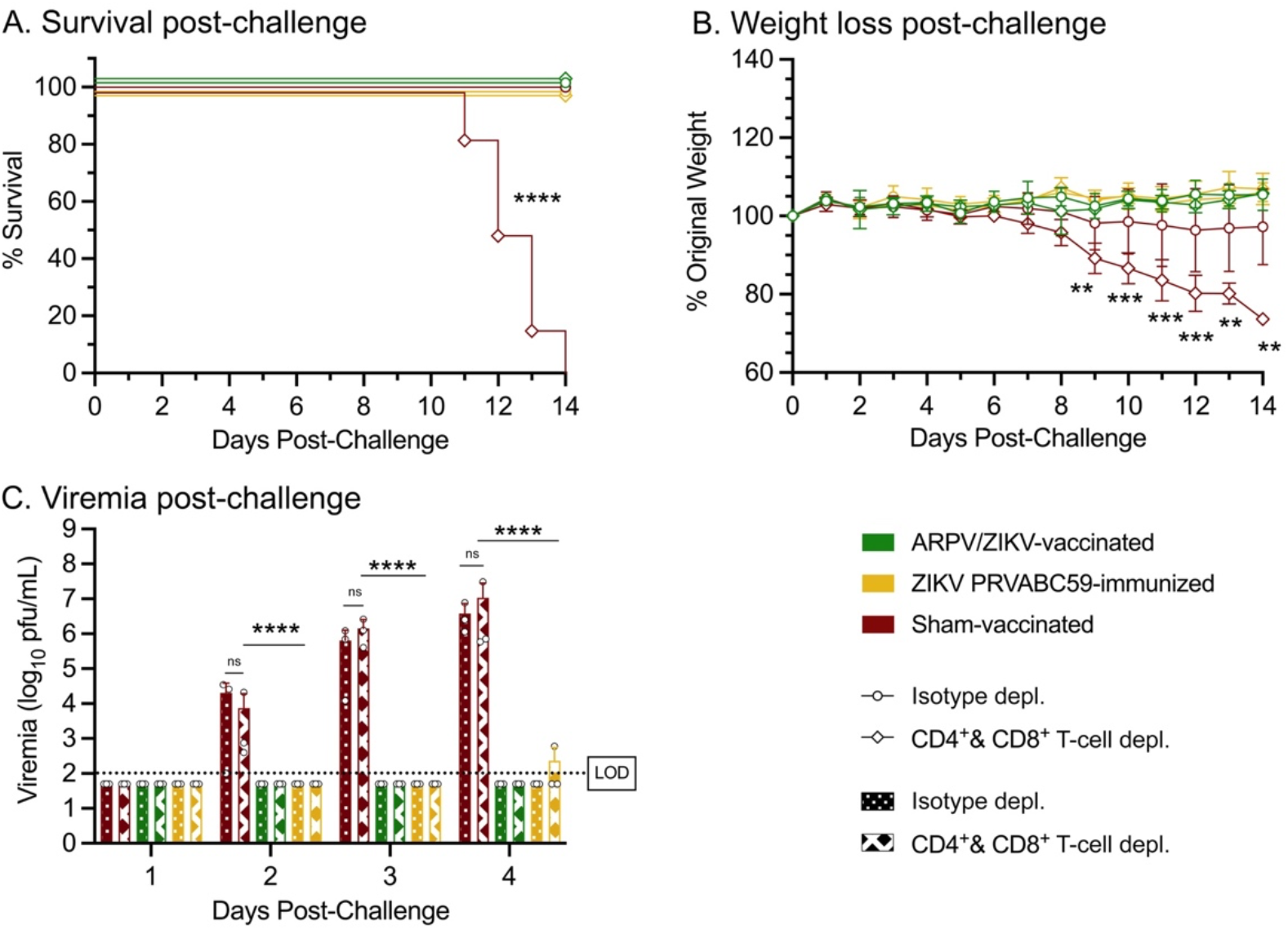
Depletion of T-cells in ARPV/ZIKV-vaccinated immunocompetent mice at challenge did not significantly impact clinical outcomes. Four-week-old C57BL/6J mice were vaccinated as previously described. Regimens of *in vivo* T-cell depletion monoclonal antibody were administered according to depletion group (anti-CD4^+^, anti-CD8^+^, both, or isotype depletion). At 30 days post-vaccination, mice were challenged with ZIKV. Mice were then monitored for (a) survival and (b) weight change post-challenge. (c) Sera was collected for 1-4 days post-challenge. LOD is 100 pfu/mL. (b) Symbols represent mean values. (c) Columns represent mean values, and symbols represent individual data points. Error bars indicate SD of the mean. Asterisks indicate significance compared to isotype depleted control mice, unless otherwise indicated: not significant (ns), *p* ≤ 0.002 (**), *p* ≤ 0.0002 (***), *p* ≤ 0.0001 (****).

To further assess the contribution of T-cells to protection during challenge, CD4^+^ and CD8^+^ T-cells were isolated from vaccinated C57BL/6J mice and transferred to naïve IFN-αβR^-/-^ mice prior to challenge. Mice that received ARPV/ZIKV-primed CD4^+^ or CD8^+^ T-cells had a statistically significant median survival time of two days longer than negative control groups that received unprimed T-cells or no T-cells (Fig. 5a-b). Mice receiving ARPV/ZIKV-primed CD8^+^ or CD4^+^ T-cells weighed more than mock transfer or no transfer mice at 4-7 dpc, or 4 and 6 dpc (respectively) (Fig. 5c-d).

**Figure 5:**
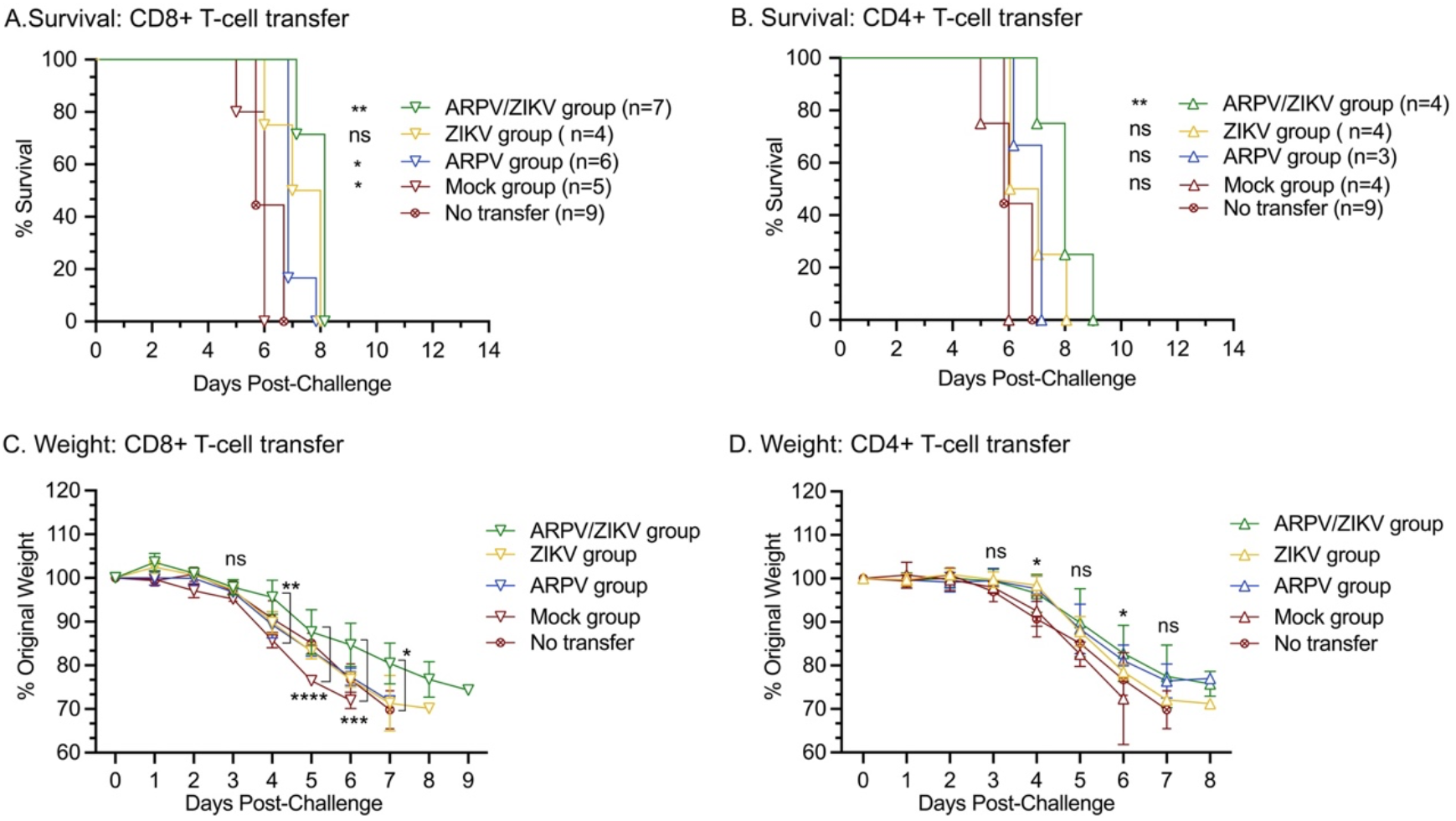
Adoptive transfer of ARPV/ZIKV-primed T-cells to naïve mice pre-challenge resulted in increased median survival times. Single-cell suspensions of CD4+ or CD8+ T-cells were prepared from the spleens of C57BL/6J mice 30 days after vaccination, then injected into naïve IFN-αβR^-/-^ mice 24 hours before challenge. Mice were then monitored for (a) survival and (b) weight change. Blood samples were taken 1 and 4 days post-challenge to quantify viremia. LOD is 100 pfu/mL. (b) Symbols represent mean values. (c) Columns represent mean values, and symbols represent individual data points. Error bars indicate SD of the mean. Asterisks indicate significance compared to the “no transfer” group, unless otherwise indicated: not significant (ns), p ≤ 0.033 (*), p ≤ 0.002 (**).

### T-cells contribute to the development of protective immune responses after ARPV/ZIKV vaccination

To determine the importance T-cell responses in developing protective immunity after vaccination, rather than their contribution to protection at the time of ZIKV challenge, Rag1 KO, Tcra KO, and muMt^−^ mice were vaccinated and challenged. By 28 dpv, no mice developed ZIKV-specific nAb titers above the limit of detection (data not shown), except for one ZIKV-immunized Tcra KO mouse which had a PRNT_50_ of 20. After ZIKV challenge, ARPV/ZIKV-vaccinated and sham-vaccinated Tcra KO mice experienced 100% mortality by 16 dpc (Fig. 6a), but surviving ZIKV-immunized Tcra KO mice began to stabilize in weight after 17 dpc (Fig. 6b). However, among muMt^−^ mice, only the sham-vaccinated group experienced significant mortality and weight loss, achieving 83% mortality by 10 dpc (Fig. 6c-d). Rag1 KO mice showed no significant differences in survival or weight loss between immunization groups (Fig. 6e-f). ZIKV-immunized Tcra KO mice and Rag1 KO mice had significantly lower viremia compared to sham-vaccinated mice on 3 dpc and 4 dpc, respectively, whereas there were no significant differences in viremia between immunization groups in muMt^−^ mice for 1-4 dpc (Fig. 6g-j). Results depicted in Figure 6 were also analyzed and presented according to immunization group in Fig. S5.

**Figure 6:**
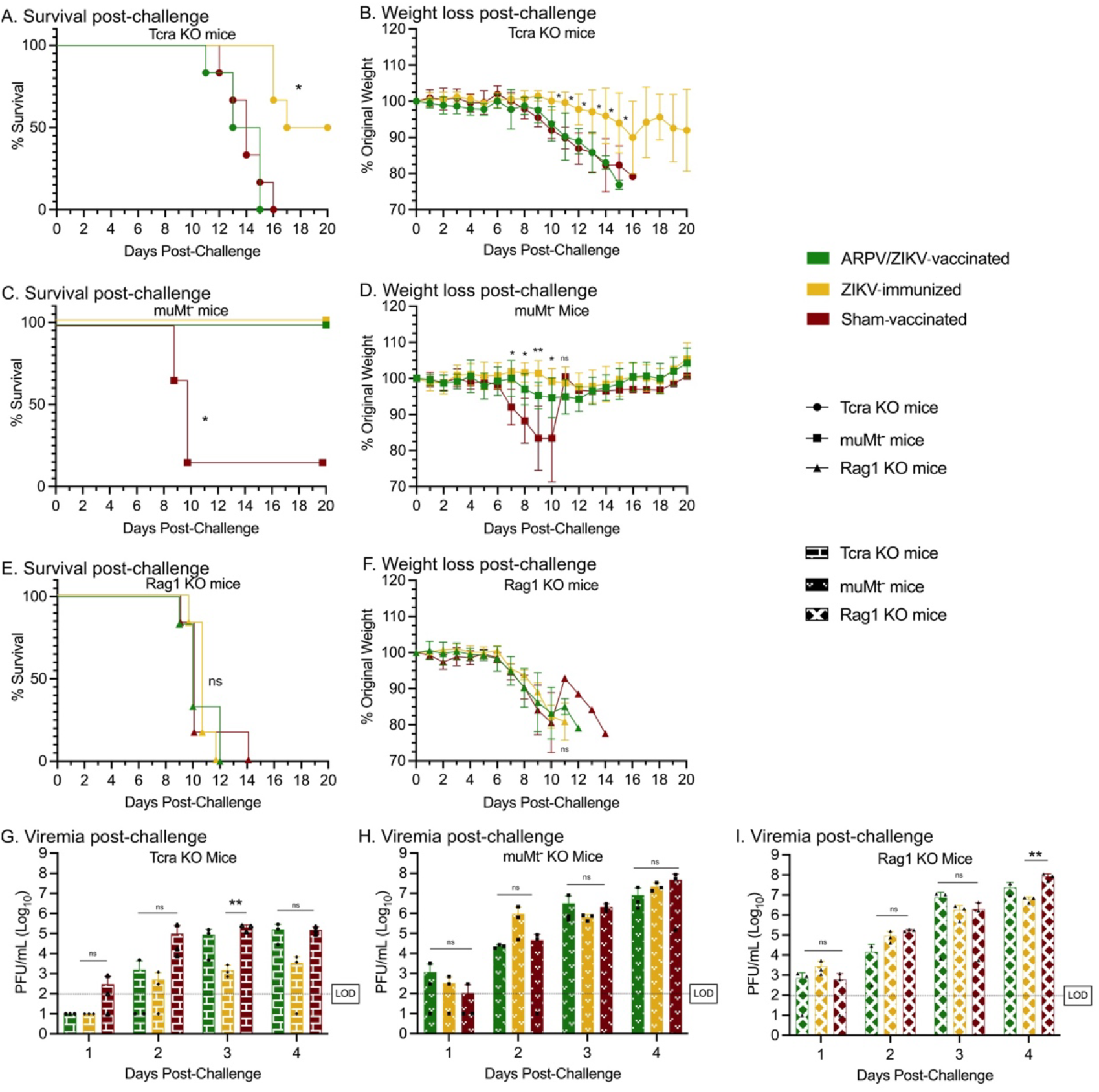
T-cells are critical for the development of an effective immune response to ARPV/ZIKV vaccination. Four-week-old mice with deficient B-cell responses (muMt^−^), deficient T-cell responses (Tcra KO), or both (Rag1 KO) were vaccinated as previously described. At 30 days post-vaccination, all mice were challenged with ZIKV. Tcre KO mice (a-b), muMt^−^ mice (c-d), and Rag1 KO mice (e-f) were then monitored for survival and weight change. (g-i) Sera was collected for 1-4 days post-challenge to quantify viremia. Dotted lines indicate the 100 pfu/mL limit of detection (LOD). (b, d, f) Symbols represent mean values. (g-i) Columns represent mean values, and symbols represent individual data points. Error bars indicate SD of the mean. Asterisks indicate significance compared to sham-vaccinated control mice, unless otherwise indicated: not significant (ns), *p* ≤ 0.033 (*), *p* ≤ 0.002 (**).

## 4. Discussion

ISFV-based vaccines may offer good alternatives to traditional flavivirus vaccine platforms due to their high degree of safety, and may be useful with patients for which live vaccines are contraindicated. Certain ISFV-based vaccine candidates also have single-dose efficacy [9,14,15], as opposed to many traditional killed or subunit vaccine candidates [28]. This is likely at least partially due to efficient genome delivery to target cells for pattern recognition receptor (PRR) detection since ARPV/ZIKV infection of mammalian cells robustly induced PRR, B-cell receptor, and antigen presentation signaling pathways, as well as T_h_1/T_h_2 T-cell polarization, which is in agreeance with other studies that indicate T_h_1 responses to be the primary ZIKV CD4^+^ T-cell responses [21]. This study also helps confirm the replication defective nature of the vaccine since there were no differences in host gene expression between infection with UV-inactivated and live ARPV/ZIKV.

Previous studies show that ISFV-based vaccines can induce exceptionally high nAb titers [9,14]. However, the underlying mechanisms of immune induction by this new type of vaccine platform, particularly regarding the role of CD4^+^ or CD8^+^ T-cell responses, remain largely unknown. For flaviviruses, neutralizing antibodies are critical for mediating protection [17,18], and cytotoxic CD8^+^ T-cells have been shown to aid in protection against ZIKV [18,20-22]. During ZIKV infection, CD4^+^ helper T-cells promote the development of nAb responses [19,29], perform cytotoxic functions and direct the immune response through cytokine-secretion [19], and may potentially protect against nervous tissue damage or provide limited protection during pregnancy [19,20]. Here, we explored T-cell-mediated responses to ISFV-based vaccines using ARPV/ZIKV as a model.

During the passive transfer study, mice passively immunized with ARPV/ZIKV-induced immune sera were incompletely protected after challenge, despite having high nAb titers. This observation, alongside previous *ex vivo* and *in vitro* T-cell and cytokine response data [9], led to the hypothesis that ARPV/ZIKV-induced cellular immunity contributes to host protection during ZIKV challenge. However, ARPV/ZIKV-vaccinated mice that were depleted of T-cells prior to ZIKV challenge did not experience significantly increased morbidity or mortality. Additionally, adoptive transfer of ARPV/ZIKV-primed CD4^+^ and CD8^+^ T-cells to naïve mice prior to challenge resulted in only a two day increase in median survival times compared to controls. Hence, ARPV/ZIKV-induced T-cell responses are partially protective during challenge, but T-cell depletion and adoptive transfer studies indicate they may play minor role and be somewhat redundant compared to the humoral immune responses. If this is the case, the partial protection after passive immunization with ARPV/ZIKV-immune sera might be due to ARPV/ZIKV’s inability to induce antibodies against ZIKV proteins other than prM and E, such as ZIKV capsid or secreted nonstructural protein 1 (NS1). NS1-induced antibodies may provide some protection against disease, as shown previously for DENV [30]. Further studies are warranted to characterize the complete antibody repertoire generated by ARPV/ZIKV vaccination compared to ZIKV infection, identify target epitopes, and explore the role of non-neutralizing antibodies in protection from disease. Overall, these results also indicate that ISFV-based vaccine platforms may benefit from adjuvants or optimization strategies to increase T-cell responses that are protective independent of their antibody helper functions.

In addition to investigating the role of T-cells in mediating protection during challenge, we also began to examine the importance of T-cells in the context of developing ARPV/ZIKV-induced immune responses post-vaccination. ARPV/ZIKV vaccination of T-cell-deficient Tcra KO mice resulted in significantly worse clinical outcomes post-challenge compared to ZIKV immunization. Hence, T-cells likely play a critical role in developing immunity after primary exposure to ARPV/ZIKV. T-cell responses were sufficient for protection in both ARPV/ZIKV- and ZIKV-immunized muMt^−^ mice, despite their lack of mature B-cells and subsequent absence of nAbs. However, this survival of ARPV/ZIKV-vaccinated muMt^−^ compared to Tcra KO mice does not necessarily indicate that T-cell reactions are more important than B-cell reactions since mice deficient in T-cells likely lacked significant T-cell-dependent B-cell activation. Future investigation will further explore the underlying mechanisms. Overall, these studies are consistent with previous ZIKV studies using similar mouse models [31].

Interestingly, ARPV/ZIKV-vaccinated muMt^−^ mice had significantly higher viremia 4 dpc than vaccinated T-cell-deficient mice, despite Tcra KO mice experiencing significantly more weight loss and death. It could be possible that Tcra KO mice had some T-cell independent B-cell activation which may have helped to control viremia during early infection. However, during late infection, B-cell-deficient mice may have been better protected due to their fully intact T-cell responses. In general, T-cell responses are able to combat infection even after neurotropic flaviviruses gain access to immune-privileged regions [17]. However, further studies are need to confirm this hypothesis.

Knowledge of the correlates of protection and immune responses induced by ISFV-based vaccines will aid antigen design and adjuvant selection, increase vaccine efficacy, and potentially alleviate concerns of ADE. The data shown here demonstrates that both nAbs and T-cell responses mediate the robust protection observed, with nAbs playing a larger role at the time of challenge but T-cells playing a significant role in the development of protective immunity after ARPV/ZIKV vaccination in mice. Overall, ISFV vaccine platforms are being continually refined, and it seems likely that they will prove themselves to be an essential tool for reducing the global burden of flavivirus disease

## 5. Acknowledgements

The authors thank Dr. Nisha Duggal (Virginia Tech, Blacksburg, VA, USA) for kindly providing Zika virus strains PRVABC59 and DakAr D 41524. The authors would also like to thank Melissa Makris for technical support of the flow cytometry experiments.

