## Supplemental Figure 1 for "Humoral and T-cell-mediated responses to a pre-clinical Zika vaccine candidate that utilizes a unique insect-specific flavivirus platform"

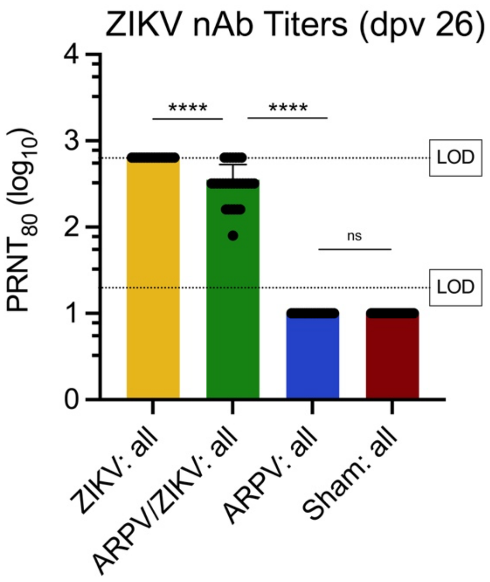


**Figure S1: Neutralizing antibodies of IFN-αβR$^{-/-}$ mice prior to T-cell depletion.** Mice were vaccinated as previously described and bled 26 days later to quantify the amount of ZIKV-specific neutralizing antibodies (nAb) in the blood. Dotted lines indicate the 20-fold and 640-fold limits of detection (LOD). Columns represent mean values, and symbols represent individual data points. Error bars indicate SD of the mean. Significance: not significant (ns), $p \leq 0.0001$ (****).
