## Supplemental Figure 2 for "Humoral and T-cell-mediated responses to a pre-clinical Zika vaccine candidate that utilizes a unique insect-specific flavivirus platform"

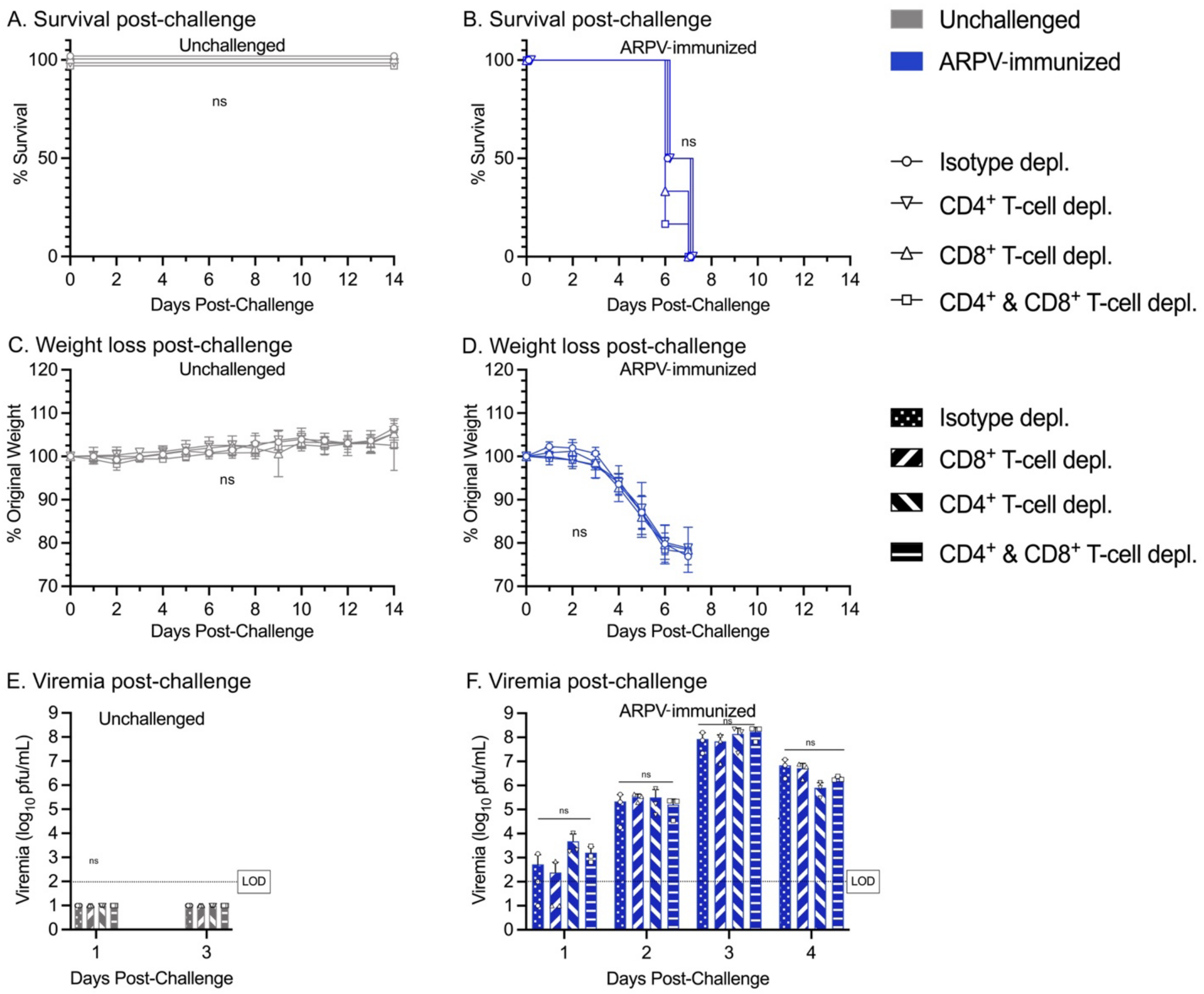


**Supplemental Figure S2: Data from unchallenged mice and ARPV-immunized mice described in Figure 3.** (a-b) Survival post-challenge. (c-d) Weight lost post-challenge. (e-f) Viremia for 1-4 days post-challenge. Dotted lines indicate the 100 pfu/mL limit of detection (LOD). (c-d) Symbols represent mean values. (e-f) Columns represent mean values, and symbols represent individual data points. Error bars indicate SD of the mean. Not significant (ns).
