## Supplemental Figure 3 for "Humoral and T-cell-mediated responses to a pre-clinical Zika vaccine candidate that utilizes a unique insect-specific flavivirus platform"

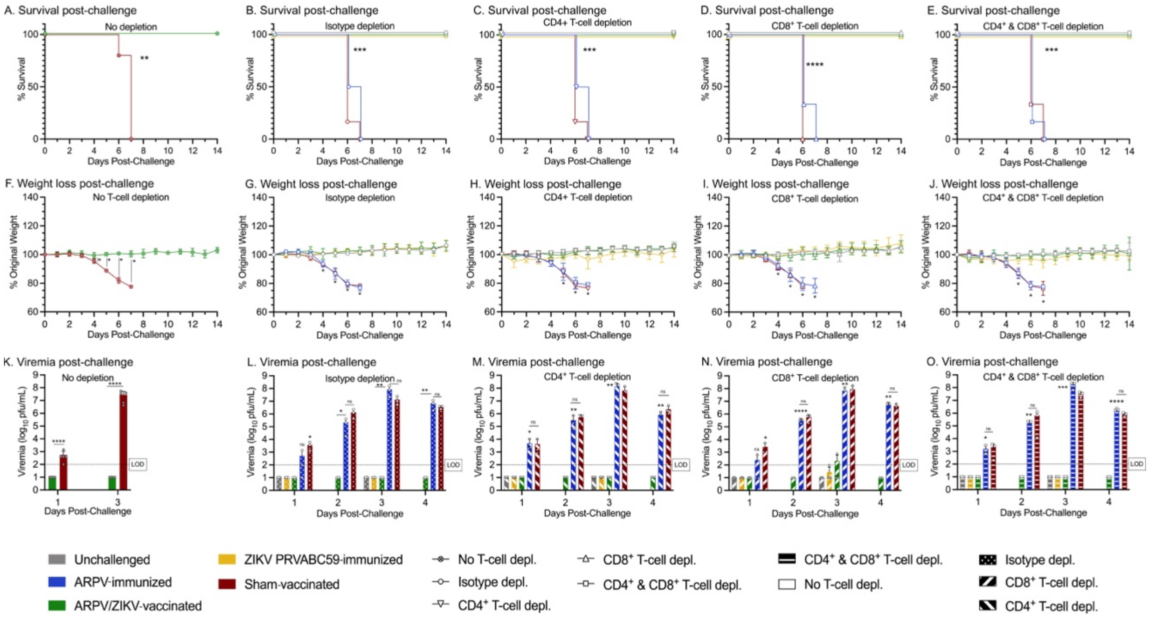


**Supplemental Figure S3: Data from the experiment described in Figure 3 was re-analyzed according to depletion group rather than immunization group.** (a-d) Survival post-challenge. (f-j) Weight loss post-challenge. (k-o) Viremia post-challenge. Dotted lines indicate the 100 pfu/mL limit of detection (LOD). (f-j) Symbols represent mean values. (k-o) Columns represent mean values, and symbols represent individual data points. Error bars indicate SD of the mean. Asterisks indicate significance compared to healthy mice (unchallenged controls), unless otherwise indicated: not significant (ns), $p \leq 0.033$ (*), $p \leq 0.002$ (**), $p \leq 0.0002$ (***), $p \leq 0.0001$ (****).
