## Supplemental Figure 4 for "Humoral and T-cell-mediated responses to a pre-clinical Zika vaccine candidate that utilizes a unique insect-specific flavivirus platform"

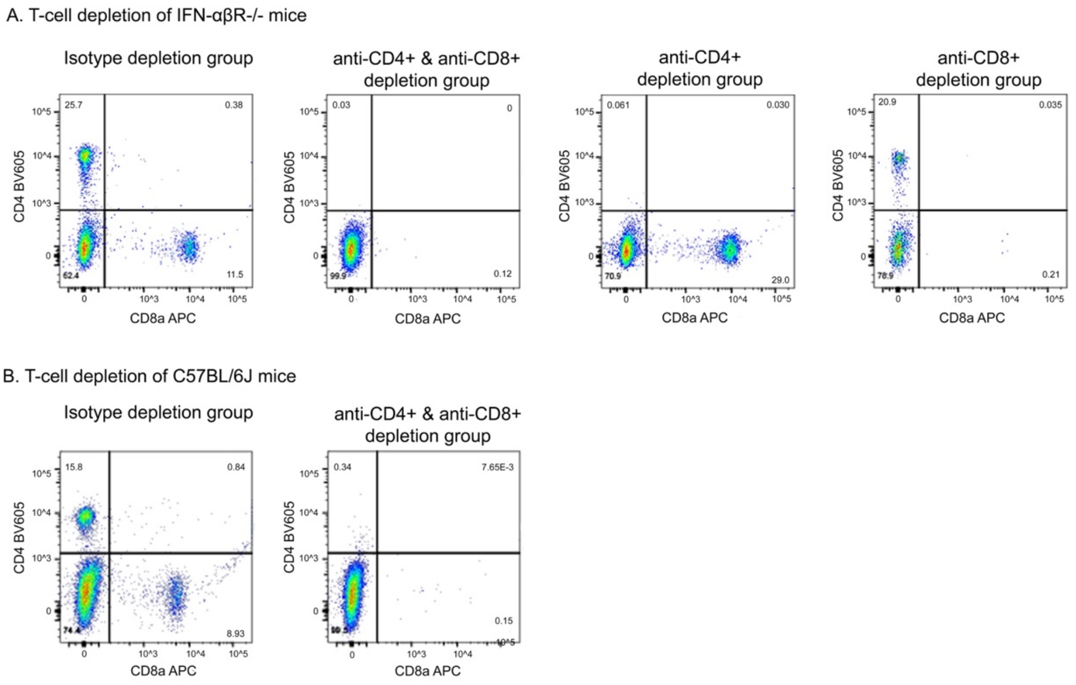


**Supplemental Figure S4: Confirmation of T-cell depletion.** Groups of age-matched mice received the same T-cell depleting antibodies as experimental groups for T-cell depletion studies in (a) IFN-αβR$^{-/-}$ mice and (b) C57BL/6J mice. At 0 days post-challenge, these age-matched groups were euthanized and their blood analyzed for circulating CD4+ or CD8+ T-cells using flow cytometry to confirm efficient depletion.
