## Supplemental Figure 5 for "Humoral and T-cell-mediated responses to a pre-clinical Zika vaccine candidate that utilizes a unique insect-specific flavivirus platform"

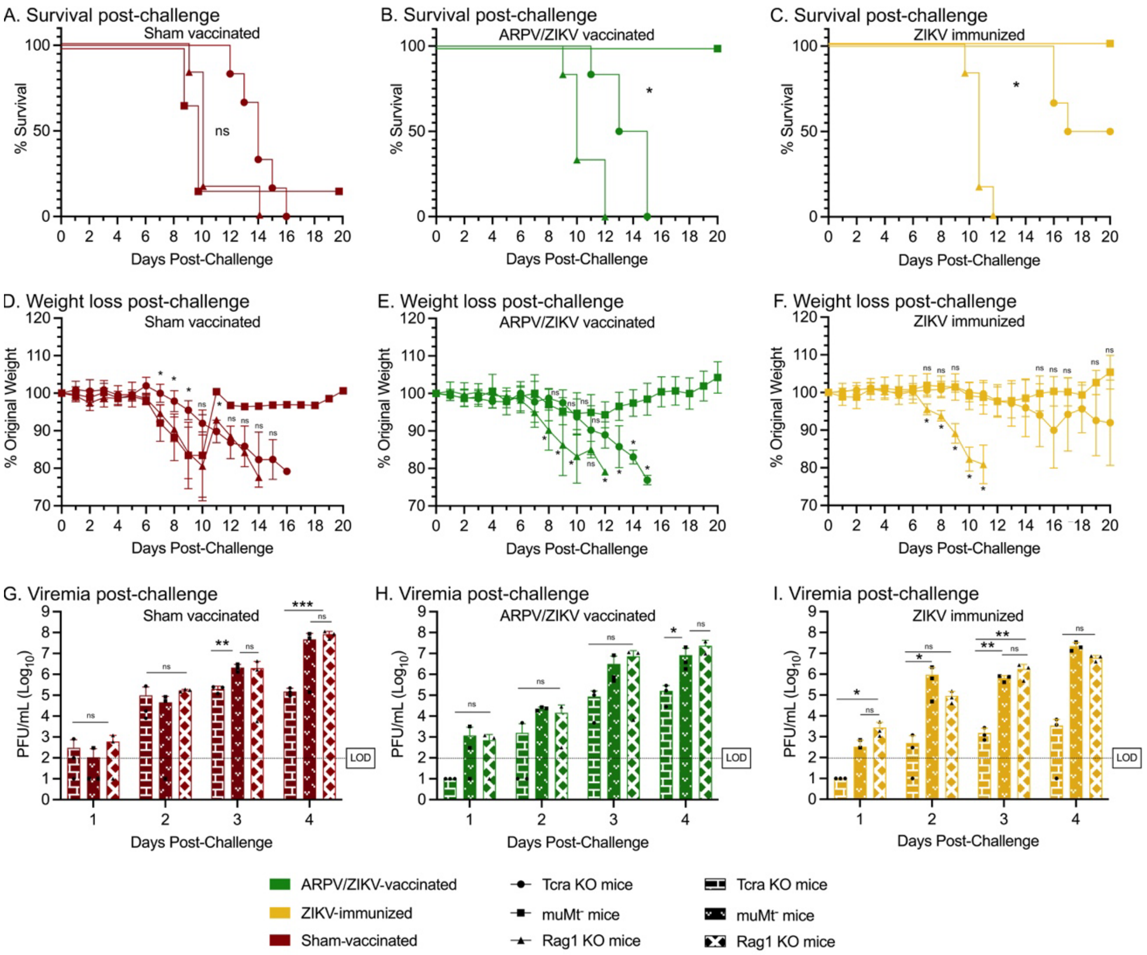


**Supplemental Figure S5: Data from the experiment described in Figure 6 was re-analyzed according to immunization group.** (a-c) Survival post-challenge. (d-f) Weight loss post-challenge. (g-i) Viremia post-challenge. Dotted lines indicate the 100 pfu/mL limit of detection (LOD). (d-f) Symbols represent mean values. (g-i) Columns represent mean values, and symbols represent individual data points. Error bars indicate SD of the mean. Asterisks indicate significance compared to muMt$^-$ mice, unless otherwise indicated: not significant (ns), $p \leq 0.033$ (*), $p \leq 0.002$ (**), $p \leq 0.0002$ (***).
